# Wound healing and anti-inflammatory potential of *Ajuga bracteosa*-conjugated silver nanoparticles in Balb/c mice

**DOI:** 10.1101/2022.09.21.508872

**Authors:** Saiqa Andleeb, Sadia Nazer, Suliman Yousef Alomar, Naushad Ahmad, Imran Khan, Abida Raza, Uzma Azeem Awan, Sadaf Azad Raja

## Abstract

**Background:** Wound therapy is complicated, uncomfortable for the patient, and costly for the health-care system. Silver nanoparticles (AgNP) have antibacterial characteristics that can prevent bacterial infection in wounds and speed up wound healing

**Objective:** The aim of current research was to investigate the wound healing and anti-inflammatory potential of biogenic synthesized silver nanoparticles (ABAgNP) using *Ajuga bracteosa* (ABaqu) in Swiss albino mice.

**Methods:** *In vivo* wound healing and anti-inflammatory activities were carried out using Bala/c mice. For *in vivo* screening of 200 mg/kg and 400 mg/kg of both ABAgNPs and ABaqu were used. Liver and kidney functional markers, hematology, and histopathological studies were carried out after 14 days of administration.

**Results:** The obtained biogenic nanoparticles were characterized, dermal toxicity, wound excision repairing, and formalin-induced paw edema assays were performed in Swiss albino mice. Dermal toxicity showed that tested concentrations of ABaqu and ABAgNPs were safe. No adverse effects, changes, and alteration in the skin of treatment groups as well as the control vehicle group (petroleum jelly) were recorded. Results revealed that the enhanced wound contraction was observed in ABaqu, ABAgNP, and the Nitrofuranose treated groups from 7^th^ to 11^th^days. The anti-inflammatory activity in formalin-induced paw edema model illustrated the potential use of silver nanoparticles ABAgNPs and ABaqu as a reducing or inflammation inhibiting agents due to the release of acute inflammatory mediators.

**Conclusion:** Therefore, it was concluded that both silver nanoparticles (ABAgNP) and *Ajuga bracteosa* (ABaqu) extracts could be used as a wound healing and anti-inflammatory agents.

## 1. INTRODUCTION

Wounds are physical injuries and commonly defined as “loss or breaking of cellular and anatomic or functional continuity of living tissues” [1, 2]. There are two conditions related to wound such as chronic or acute wound [3, 4]. Chronic wounds occur due to medical issues and results in socioeconomic crisis, disability, morbidity, and mortality of the large population [5]. During an injury, a blood clot is formed due to the damage in capillary and followed by the early phase of the inflammation. The mechanism of tissue regeneration completed in four main programmed phases that are hemostasis, inflammation, proliferation, and the maturation or remodeling [6]. Immune response to injury/wound or infection causes inflammation, results in the removal of some factors, physiological function, and tissue structure restoration [7]. Two types of drugs are used for the treatment of inflammation non-steroidal drugs and steroidal drugs. On the other hand, these types of drugs cause various side effects, so, it is the need to find alternative medicines that can solve the problems efficiently, well-tolerated, homeostatic, and modulatory *via* the body [8].

Infections related to various wounds, elevate medical expenditures and put a strain on health-care systems due to prolonged hospital admissions, treatment failures, infection persistence, and delayed wound healing, which often leads to amputation and increased death [9]. New antibiotics and antibacterial techniques are urgently needed to tackle infections in wounds. Chemically manufactured medications have several side effects; as a result, medicinal plants are being studied for their curative potential due to their widespread availability and lack of toxicity for therapeutic usage while being effective at preventing infections in different wounds [10]. Many researchers reported the use herbal or traditional medicines for solving health issues [5, 11]. Plants or synthesis of medicine from the plant is considered an important source of recent medicines worldwide [12]. In most developing countries (like Ethiopia), the treatment of dermatological problems as well as skin disorders like wounds, burns, and cuts depend upon the use of medicinal plants that play the main role in wound healing [13-15, 2]. Products isolated from plants have been used as an effective anti-inflammatory agent having low toxicity and effective pharmacological property [16]. *Ajuga bracteosa* is a medicinal herb that grows from Azad Kashmir to Nepal at an elevation of 2000 meters in the sub-Himalayan tract. Its leaves are used to treat several diseases like stomach acidity, burns, boils, wound healing, hypertension, jaundice, and sore throats. It has also been demonstrated to have anti-inflammatory, anticancer, antimicrobial and antimalarial properties [17].

The best new alternative and effective procedure for the new drugs development and drug delivery related to inflammation is nanotechnology. It is the technique that is used to synthesize the product at a nano-level about 1-100 nm [18]. Metal nanoparticles and their effectiveness in fighting infectious diseases is an attractive area of research with many applications [19]. There are various nanoparticles but because of the potent antimicrobial properties against bacteria, viruses, and fungus, silver nanoparticles have been utilized for over a century. Furthermore, silver’s anti-inflammatory properties as well as its ability to speed wound healing have recently been demonstrated [20]. Phyto-nanotechnology, in which nanoparticles are manufactured utilizing extracts from plants has attracted a lot of attention as a simple, quick, scalable, and cost-effective alternative. Plant extracts have the benefit of biocompatibility since they are high in bioactive chemicals that are amenable to extraction using water as an inert solvent and may also be used as reducing and capping agents in the creation of nanoparticles [21]. The inherent property of antibacterial suppression in antibiotic resistant bacteria has been reported in silver nanoparticles produced from plants that are widespread in secondary metabolites [22, 23]. Keeping in mind the importance of phytochemical assisted biogenic synthesis technique and medicinal significance of *A. bracteosa*, here we report the anti-inflammatory and wound healing effect of the synthesized biogenic silver nanoparticles using *A. bracteosa* (ABAgNP).

## 2. MATERIALS AND METHOD

### 2.1 Ethical statement

All experiments were appropriately performed in accordance with the Institutional Review Board (IRB) bioethics approval. The experiments have been designed to avoid distress, unnecessary pain and suffering to the experimental animals. All procedures were conducted in accordance with international regulations referred as Wet op de dierproeven (Article 9) of Dutch Law. Ethical approval has been provided by Office of Research Innovation and Commercialization (ORIC), University of Azad Jammu and Kashmir, Muzaffarabad, Pakistan vide No. 649/ORIC/2021; Dated: 16-12-2021.

### 2.2 Chemicals used

All chemicals and reagents were obtained from Sigma Aldrich (Germany), Merck (Germany), and Sigma Aldrich (Switerzland). silver nitrate (AgNO_3_), dimethyl sulphoxide (DMSO), formalin, petroleum jelly, nitrofuranose, diethyl ether, formaldehyde, Diclofanic sodium.

### 2.3 Extract preparation, silver nanoparticles synthesis, and characterization

The roots and aerial parts of *A. bracteosa* were cleaned with the running tap water to remove dust and air-dried at room temperature (20 ºC±2) under shade. The dried material was crushed in to fine powder and used for the aqueous extract preparation *via* maceration [24]. Silver nanoparticles using ABaqu were synthesized by simple ratio method [25]. Briefly, aqueous extract was prepared by adding 10 g of root or aerial part powder to 100 mL of distil water (1:10 ratio) and heated in a water bath at 50°C for 40 min. The solution was then allowed to cool at room temperature (RT, 25°C) followed by filtration through Whatman filter paper (No. 1). 5 mL of this prepared extract was slowly added into 45 mL of silver nitrate (AgNO_3_, 1 mM) in an Erlenmeyer flask. The resultant mixture was adjusted to pH 7 and incubated on a rotary shaker at 200 rpm, in dark, for 24 h at RT. During this period, visual observations were made to detect any change in color. The synthesized nanoparticle suspension was centrifuged at 4,500 rpm for 20 min to collect ABAgNPs as a pellet. To ensure removal of unreacted silver ions and any unbound phyto-constituent, this pellet was washed thrice with distil water, air-dried, and stored at room temperature for further use. The preliminary characterization of green synthesized nanoparticles (ABAgNPs) was done through a UV-viz spectrophotometer (between 200 nm to 800 nm). The morphology (shape and size) of ABAgNPs and ABaqu were confirmed through a scanning electron microscope (SEM). Fourier transform infrared spectroscopy (FTIR, Thermo Fischer Scientific) recorded the absorption spectra of dried pellets of ABaqu and ABAgNPs in the range of 4,000–400 cm^1^ to identify the presence of functional groups involved in bio-reduction.

### 2.4 Experimental animal

The male Swiss albino mice (23-30 g) were purchased from the animal facility center of the National Institute of Health (NIH), Islamabad, Pakistan. The animals were acclimatized for 2 weeks before the beginning of the current research. Animals were maintained in individual cages with free access to food and water under controlled conditions (5 % humidity, 12 h day/night cycles, and the room temperature at 25±2 °C). The handling and special care of the Swiss albino mice were according to the established public health guidelines in Guide for Care and Use of Laboratory Animals [26]. The feed (commercial chow) and water consumption patterns were observed throughout the study period.

### 2.5 Acute Dermal Toxicity

Acute dermal toxicity study was performed with a limit test at 1000 mg/kg (1 ml) as a single dose for both ABaqu and ABAgNPs according to the adopted procedures described previously with slight modifications [27]. The mice were selected randomly after acclimatization to the laboratory conditions, and kept in their cages for at least 5 days before dosing. Three groups having three healthy Balb C mice weighing 23–30 g were acclimatized such as group I treated with vehicle (petroleum jelly); group II animals treated with ABaqu: and group III animals treated with ABAgNPs. For acute dermal toxicity, the fur was removed 24 h before the study from the dorsal area of the trunk of the tested animals and the highest concentration of the ABaqu and ABAgNPs (1000 mg/kg) ointment 10% (w/w) for a period of 24 h were applied. The treatment was repeated for 14 days. After application, cage side observation was performed daily for the next 14 days to notice the late development of dermal toxicity [28]. Tissues from all the experimental treatments were collected at the end of the experiment by using sterile scissors and forceps for histopathological studies. The tissue samples were fixed in 10% formaldehyde for 1 week for the histological examination [29]. Water and food were provided to all animals regularly ad libitum.

### 2.6 Excision wound model

The wound-healing effect of ABAgNPs and ABaqu were evaluated in Swiss albino mice using the protocol of Sasidharan *et al*. [30] with slight modifications. Mice were anesthetized using diethyl ether (45 mg/kg) given by the intraperitoneal route. Fur was removed from the dorsal area and a full-thickness wound of approximately 1.5 cm x 1.5 cm was made on the shaved area. Six groups (n=6) were made for the wound healing evaluation; group I had no wound and used as normal control, group II wound was treated with vehicle (petroleum jelly), group III was not treated and left open, group IV was treated with nitrofuranose ointment in petroleum jelly, group V was treated with *A. bractoesa* extract (ABaqu; 400 mg/kg), group VI was treated with synthesized silver nanoparticles (ABAgNPs; 400 mg/kg) using *A. bractoesa*. The decrease in wound diameters during the healing process was measured with the help of scale in millimeters. The wounded mice were kept for 14 days for the treatment and further observations. Tissues from mice wounds were collected at the end of the experiment by using sterile scissors and forceps for histopathological studies. The tissue samples were fixed in 10% formaldehyde for 1 week for the histological examination [29].

### 2.7 Formalin-induced paw edema

*In vivo* anti-inflammatory effect of *A. bractoesa* aqueous extract ABaqu (400 mg/kg), green synthesized ABAgNPs using *A. bractoesa* (400 mg/kg) were screened in Swiss albino mice (Balb/c) 31. The experiment was designed for 11 days. Swiss albino mice were divided into 6 groups (n=6) such as group I (Normal control), group II (vehicle; dH2O), group III (2% formalin; 0.1 ml), group IV (2% formalin + Diclofenac sodium), group V (2% formalin + ABaqu; 400 mg/kg), and group VI (2% formalin + ABAgNps; 400 mg/kg), and treated daily accordingly. Paw edema was induced by injection of 2% formalin (0.1 ml) into right hind paw of the rats [32]. The volume of formalin-induced paw edema was measured with Vernier-calliper after each hour up to 11^th^ day, and compared with control during the experiment. At the end of the experiment (after 11^th^ day) animals were sacrificed, blood and tissue samples were collected. The serum enzymes such as Alanine transaminase (ALT), Alkaline phosphatase (ALP), and Aspartate transaminase (AST) were determined using reported protocols (King, 1965). The hematological parameters like eosinophils, red blood cells, white blood cells, platelets, hematocrit, lymphocytes, neutrophils, monocytes, and hemoglobin were measured according to the standard procedures [33]. Tissues from mice wounds were collected at the end of the experiment by using sterile scissors and forceps for histopathological studies. The tissue samples were fixed in 10% formaldehyde for 1 week for the histological examination [29].

### 2.8 Statistical analysis

Each experiment was conducted thrice time. Statistical analyses were performed using graphpad prism for windows (version 5.03) and also used to plot graphs with error bars of standard errors of the means (SEM). ‘a’ indicates significant difference between control and diseased groups; ‘b’ indicates significant difference between control and other treatment groups; ‘c’ indicates significant difference between diseased and other treatment groups; ‘d’ indicates significant difference between vehicle and other treatment groups. “e” indicates significant difference between positive control and ABaqu and ABAGNPs treated groups. Values are mean±sem(n=6). Statistical icons: single letter (a/b/c/d=p≤0.05); double letter (aa/bb/cc/dd =p≤0.01); triple letter (aaa/bbb/ccc/ddd=p≤0.001).

## 3. RESULTS

### 3.1 Synthesis and characterization of nanoparticles

The synthesis of ABAgNPs was confirmed through the UV-Viz spectrum at 400 nm. The tube-like partcles as well as several aggregates of nanoparticles with a diameter ranging from 5 µm up to 50 µm was recorded *via* SEM. The IR spectrum had shown bands correspond to O-H and H stretching and confirmed the presence of alcohol and phenol’s, Carbonyl stretching, N-H and C-N stretching indicating the presence of aromatic amino groups and proteins. The absorption bands at 1108 cm^-1^ and 1021 cm-1 might have contributed by the C─O group of the polysaccharides.

### 3.2 Acute dermal toxicity

Results revealed that the maximum concentration of the ABaqu and ABAgNPs ointment (10% w/w) was not toxic and after 24 h, the dermal site did not show any sign of inflammation or irritation (Figure 1). No sign of toxicity was observed when the mice were followed for 48 h. Moreover, there was no mortality were detected during the 14 days. The hair growth was appeared and easily visualized in the ABaqu and ABAgNPs treated groups (Figure 1). Through histology no inflammatory cells were found in all treated groups. Not any difference was observed when ABaqu and ABAgNPs treated groups compared to vehicle treated group Normal epithelization was recorded (Figure 1). Therefore, we can say that ABaqu and ABAgNPs may be safe at 1000 mg/kg (10% w/w) and the oral LD_50_ is greater than tested dose in the Swiss albino mice. The WBC level was noted as slightly high in ABaqu treated animals and ABAgNPs treated animals (3.75±0.23 10^3^/UL and 3.57±0.47 10^3^/UL respectively) compared to vehicle treated group (3.47±0.638 10^3^/UL). Level of RBC was recorded as slightly low in ABaqu treated animals (4.59±0.19 10^6^/UL) and remains normal in ABAgNPs treated animals (4.8±0.19 10^6^/UL) when compared to vehicle group (4.83±0.5 10^6^/UL). Slight difference was noted in hemoglobin and hematocrit level of ABaqu treated animals. The hemoglobin level of ABaqu treated animals was 12.66±1.15 g/dl and remains normal in ABAgNps treated animals (13.3±0.57 g/dl) compared to vehicle control group (13.0±1.0 g/dl) and there was no significant difference noted. Hematocrit level in vehicle was noted as 38 ± 8.1 % and was slightly high in both ABaqu (40.0±3.5 %) and slightly low in ABAgNPs (37.0±4.7 %). Platelets and Neutrophil levels slightly increased in both ABaqu and ABAgNPs treated animals respectively compared to (DH_2_O) control group. Eosinophil, lymphocytes, monocytes, and MCV in both treated groups ABaqu and ABAgNPs showed no significant difference when compared to vehicle (DH_2_O).

**Fig. 1.**
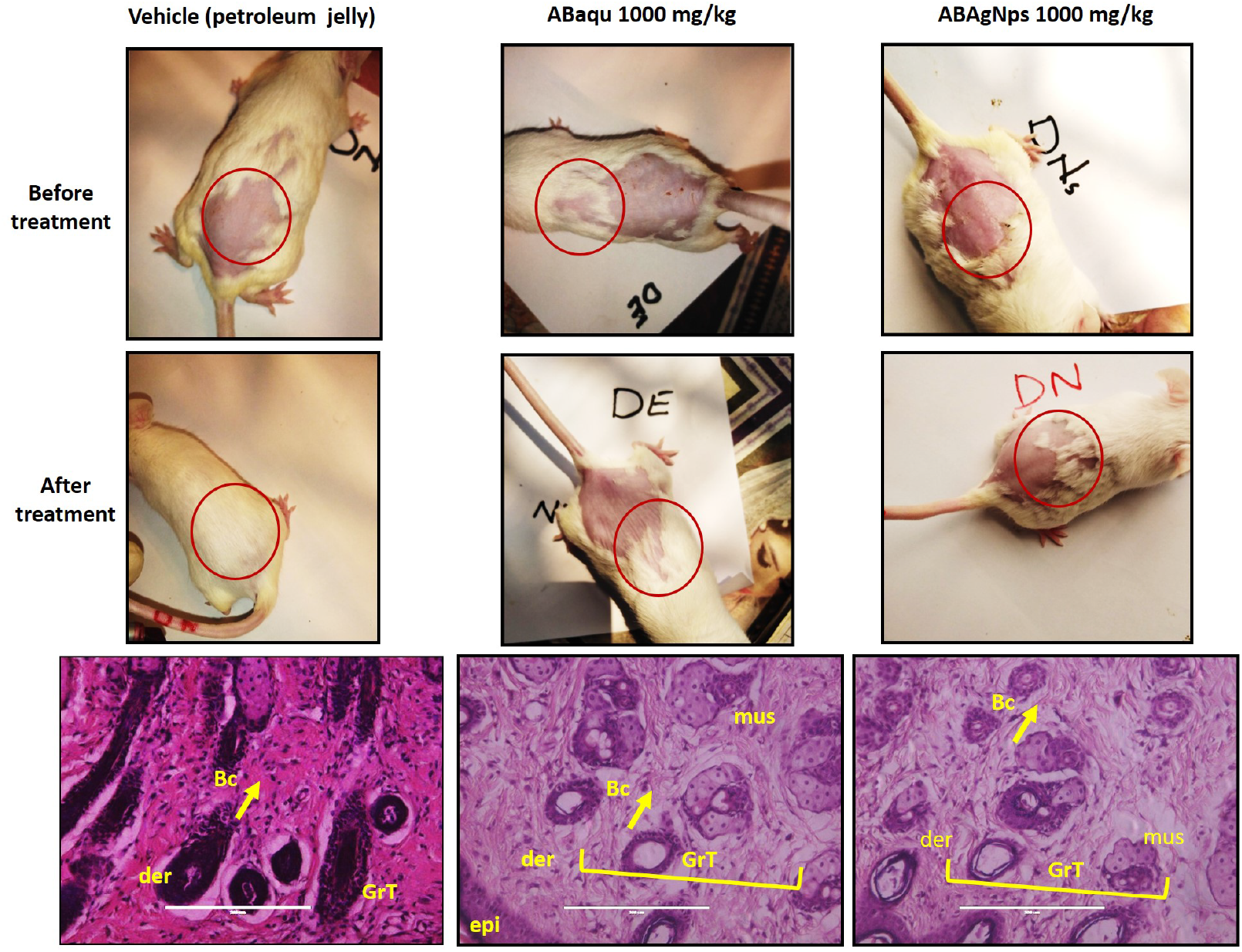
Effect of *Ajuga bracteosa* conjugated silver nanoparticles during acute dermal toxicity assay in Swiss albino mice.

### 3.3 Wound healing effect

The wound-healing activity of synthesized ABAgNPs was confirmed in Swiss albino mice using an excision wound model. The changes in the wound appearance were took place on 1^st^ day, 3^rd^, 5^th^, 7^th^, 9^th^, and 11^th^ day. On 3^rd^ day Nitrofuranose, ABaqu, and ABAgNPs treated mice exhibited brown color, moist and soft scabs while vehicle (petroleum jelly) and open wound mice showed thick, hard and intact scabs. On 11^th^ day, re-epithelialization was observed in Nitrofuranose, ABaqu, and ABAgNPs treated mice (Figure 2). The enhanced wound contraction in ABaqu, ABAgNPs, and the Nitrofuranose treated groups from 7^th^ to 11^th^ days were recorded. Both of the ointments (ABaqu and ABAgNPs) showed significant wound healing than open wound (control group). On 11^th^ day, both Nitrofuranose and ABAgNPs showed 70% wound contraction rate compared to open wound group (10%), and vehicle (10%) and ABaqu (60%) treated groups. ABAgNPs treated wounds exhibited no evidence of pus formation and bleeding during treatment, whereas control wounds revealed the remarkable inflammation. At the end of the experiment, the ABAgNPs treated wound and Nitrofuranose treated groups showed approximately maximum of wound closure (3.0±0.0 mm and 3.0±0.1 mm) compared to vehicle (9.0±0.1 mm) and ABaqu (4.0±0.0 mm) treated groups. Histopathological examination revealed diffused inflammatory cells trapped, focal inflammatory cells infiltration, and area of hemorrhages in the open wound mice in which granulated tissue was not fully formed (Figure 3). In open wounded mice damage of epidermis, dermis and granular tissues were also recorded. On the other hand, in case of vehicle treated group epithelization, keratinization, and granulation was observed. ABaqu and ABAgNPs treated wound mice showed significant increase in epithelization and remarkable fibroblast proliferation (Figure 3). No inflammation was observed in Nitrofuranose, ABaqu and ABAgNPs treated groups. Interestingly, ABAgNPs treated wound group showed complete epithelization with keratinization (Keratin covering), and fibrous connective tissue proliferation. New blood capillaries formation was also seen in Nitrofuranose, ABaqu and ABAgNPs treated groups (Figure 3).

**Fig. 2.**
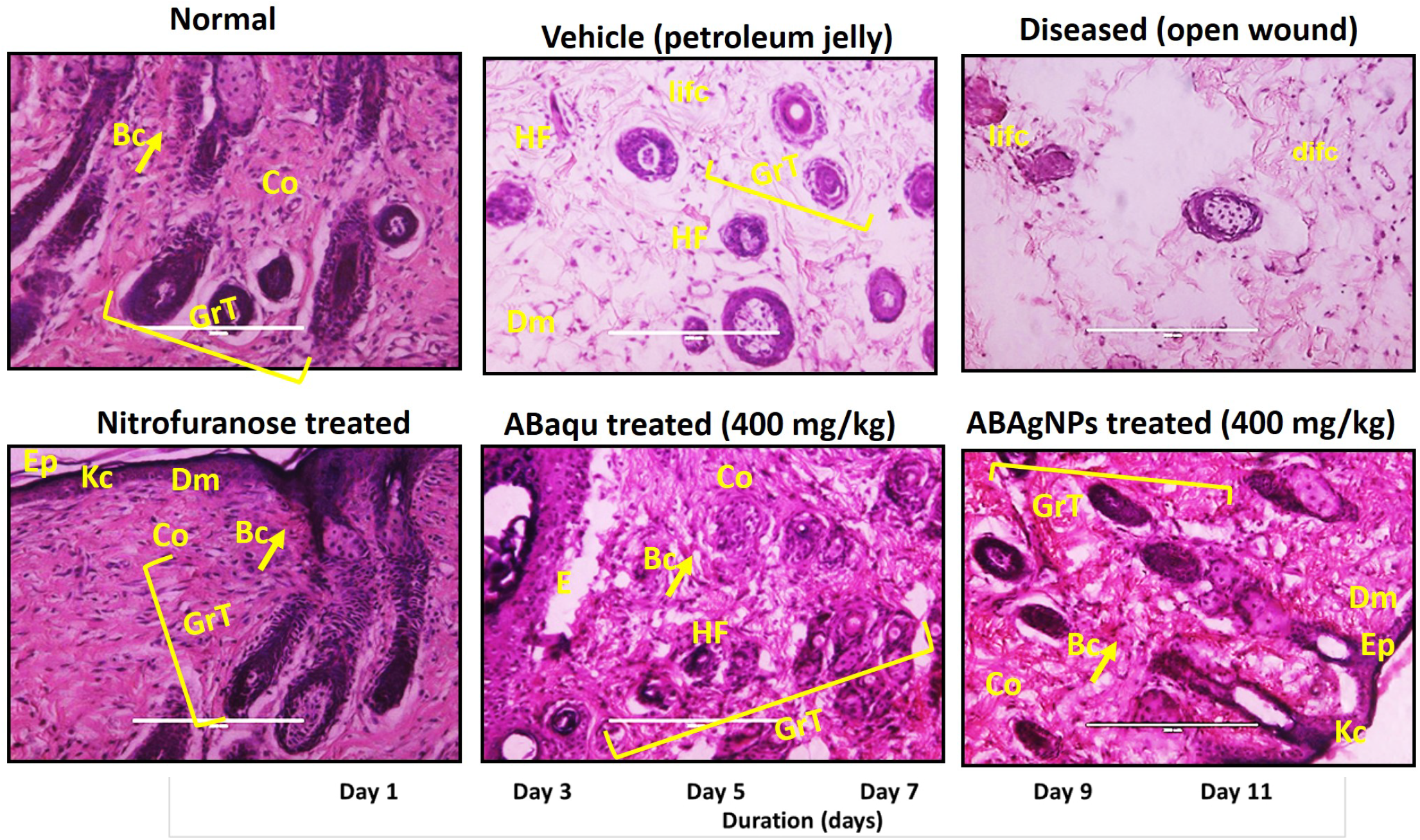
*In vivo* wound healing effect of *Ajuga bracteosa* conjugated silver nanoparticles.

**Fig. 3.** Histopathological examination of open wound and its treatment with *Ajuga bracteosa* conjugated silver nanoparticles and its aqueous extract.

### 3.4 Anti-inflammatory effect

Data revealed that formalin induced animals exhibited significant elevation of paw thickness immediately after 2 h (*P* < 0.001) of induction and regularly increased at each hour of the study period (Figure 4). Paw volume increased up to 5^th^ day. On the other hand, the aqueous extract (ABaqu) and silver nanoparticles (ABAgNPs) significantly reduced the formalin induced paw edema at 400 mg/kg tested dose (Figure 4). The standard drug diclofenac sodium also showed the anti-inflammatory activity after 5^th^ day of induction. Paw thickness was decreased in ABaqu treated formalin induced group from 0.8±0.02 mm to 0.5±0.1 mm as well as ABAgNPs formalin induced treated group (0.89±0.02 mm to 0.6±0.0 mm) compared to control and vehicle treated group (0.6±0.1 mm) which had normal paw thickness throughout the studies. Formalin induced diseased group showed increase in paw thickness (0.6±0.5 mm to 1.9±0.1 mm) after few days (Figure 4). Results revealed that both ABaqu and ABAgNPs were able to reduce the inflammation at 400 mg/kg. Figure 4 shows the histological examination of formalin induced paw edema and anti-inflammatory effect of green synthesized silver nanoparticles (ABAgNPs) and ABaqu extract. Formalin induced paw showed the focal aggregations, infiltration of diffused and focal inflammatory cells, and overcrowding of edema. These inflammations were significantly reduced by the use of ABaqu and ABAgNPs. ABaqu treated mice showed a clear keratin covering, muscular cells, well developed dermis, and normal histopathological structure. ABaqu and ABAgNPs also reduced the focal aggregations in diseased treated mice.

**Fig. 4.**
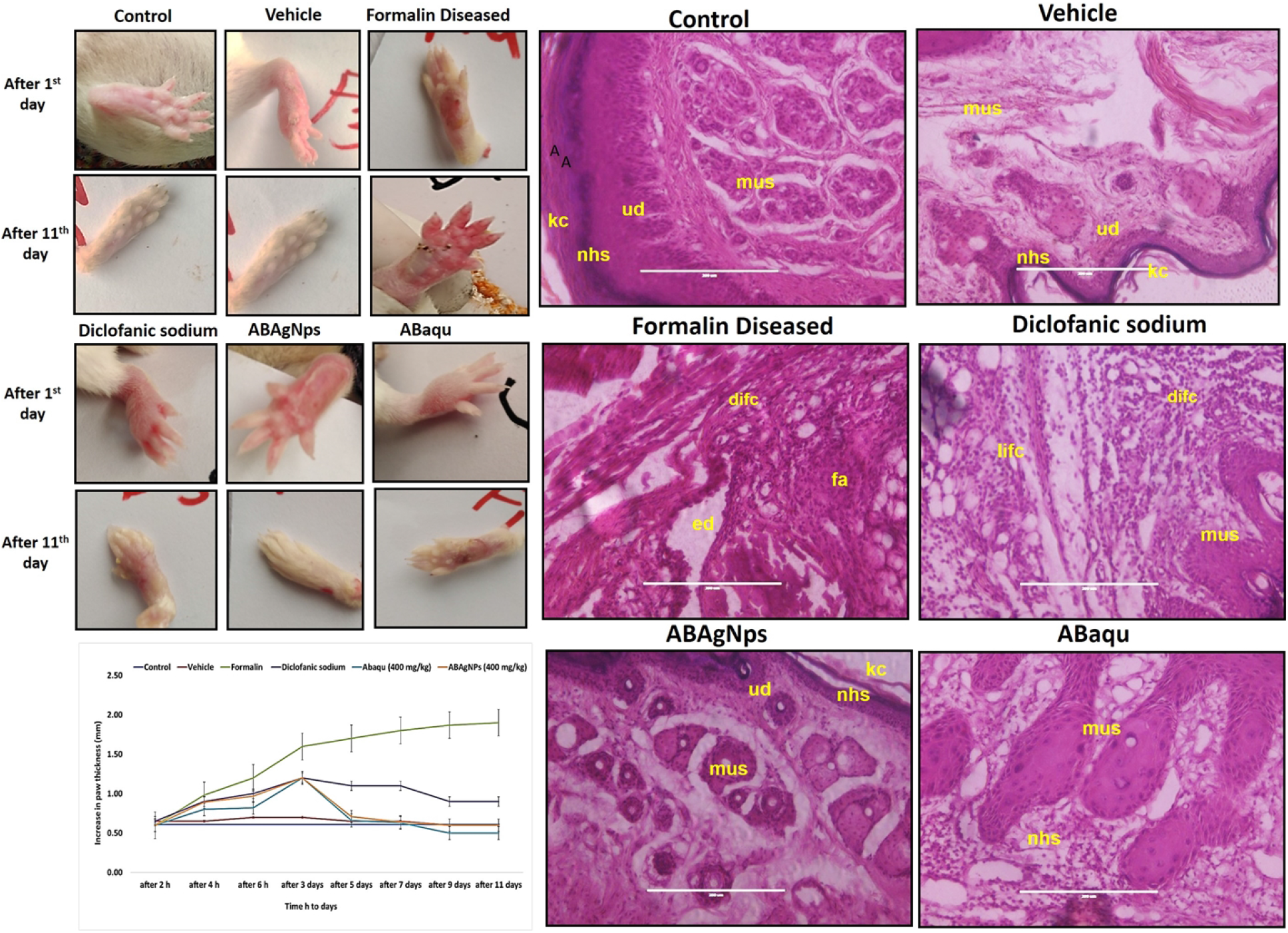
*In vivo* anti-inflammatory effect of *Ajuga bracteosa* conjugated silver nanoparticles.

The data presented in Table 1 revealed that WBC level was greater in formalin induced paw edema treated group compared to other group (8.5 ± 0.2 10^3^ µL) but it was noted that the WBC level was significantly reduced in diclofanic sodium treated group (4.2 ± 0.310^3^ µL), ABaqu and ABAgNps treated group (4.2± 0.2 10^3^ µL and 4.4± 0.4 10^3^ Ul, respectively. In control group the WBC level was noted as 3.5± 0.3 10^3^ µL. It was noted that hemoglobin and red blood cells level was high in diseased mice (10.33±0.310^6^ µL and 1.24±0.310^6^ µL) while both were increased in ABaqu (13.57±0.510^6^ µL and 4.62±0.2 10^6^ VL) and ABAgNps treated groups (13.66±0.2 10^6^ µL and 4.47±0.2 10^6^ uL) compared to others. The level of platelets was increased in formalin induced paw edema mice (837.0 ± 0.3 10^3^ µL). While the level of platelets was significantly reduced in ABaqu and ABAgNps treated group (478.33 ± 0.2 10^3^ µL and 478.0 ± 0.4 10^3^ Ul), respectively. There was increase in neutrophils level in formalin induced diseased group (38.3 ±2.7%) when compared to control group (22.3± 1.6%) as well as diclofanic sodium treated group (16.3 ±1.7%). Decreased level of neutrophils was observed in ABAgNps treated group (19.0 ± 1.9%). There was increase in lymphocytes and eosinophils level of formalin induced diseased group (86.0 ±2% and 3.0± 0.4%). Lymphocytes level was recorded as low in ABaqu (78.0± 4.3 %) and ABAgNps treated groups (76.7 ±5.6 %).

**Table 1.** Anti-inflammatory impact of green synthesized silver nanoparticles (ABAgNPs) and ABaqu extract level of total WBC, RBC, Hemoglobin, and leucocytes against formalin induced paw edema in Swiss albino mice.

| Treatments →<br>Parameter↓ | Control | Vehicle<br>DH<sub>2</sub>O | Diseased<br>(Formalin) | Diclofanic<br>sodium (DF) | ABaqu (400<br>mg/kg) | ABAgNps<br>(400 mg/kg) |
| --- | --- | --- | --- | --- | --- | --- |
| WBC 10<sup>3</sup> U/l | 3.5± 0.3 | 4.1 ±0.2 | 8.5± 0.2 | 4.2 ±0.3 | 4.2± 0.2 | 4.4± 0.4 |
| RBC 10<sup>6</sup>/U/l | 4.83±0.2 | 5.06±0.3 | 1.24± 0.3 | 3.43±0.2 | 4.62±0.2 | 4.47±0.2 |
| Hb g/dl | 13.0±0.4 | 12.4±0.3 | 10.32±0.3 | 13.42±0.4 | 13.5±0.5 | 13.66±0.2 |
| Platelets 10<sup>3</sup> U/l | 206.6± 0.3 | 257.6± 0.2 | 837± 0.3 | 795.3± 1.2 | 478.33± 0.2 | 478.0± 0.4 |
| Neutrophils % | 22.3± 1.6 | 23.3 ±0.7 | 38.3 ±2.7 | 16.3 ±1.7 | 26.3± 2.5 | 19.0 ±1.9 |
| Eosinophils % | 2.3 ±0.7 | 2.0± 0.0 | 3.0± 0.4 | 2.0±0.0 | 2.7±0.3 | 2.3±0.3 |
| Lymphocytes % | 71.3 ±3.2 | 70.7± 5.0 | 86.0 ±2 | 78.3± 2.3 | 78.0± 4.3 | 76.7 ±5.6 |

Values are mean ± SEM (n=6). p≤0.05, p≤0.01, p≤0.001

It was revealed that the ALT level elevated in formalin induced diseased mice group (121.7±10.8 U/L) compared to control groups. While the ALT level was reduced after treatment with standard drug (74.3±2.9 U/L). The significant reduction was recorded in ABaqu and ABAgNPs treated animals (95.0±4.7U/L and 76.3±3.4 U/L) as well (Table 2). It was noted that AST level elevated in formalin induced diseased mice group (112.7 ±4.2 U/L) compared to control groups. Results revealed that ABAgNPs treated mice showed significant decrease in AST level (71.3 ±4.5 U/L). ALP level found increased in formalin induced diseased mice group (280.0±11 U/L) compared to control groups. ABAqu and ABAgNps treated groups showed significant decreased in ALP level (194.3±8.4 U/L and 175.7±6.3 U/L) as shown in Table 2.

**Table 2.** Anti-inflammatory impact of green synthesized silver nanoparticles (ABAgNPs) and ABaqu extract on serum enzymes against formalin induced paw edema in Swiss albino mice.

| Treatments →<br>Parameter↓ | Control | Vehicle<br>DH<sub>2</sub>O | Diseased<br>(Formalin) | Diclofanic<br>sodium<br>(DF) | ABaqu (400<br>mg/kg) | ABAgNps<br>(400<br>mg/kg) |
| --- | --- | --- | --- | --- | --- | --- |
| ALT U/L | 46.0 ±4.1 | 59.3 ±2.5 | 121.7±10.8 | 74.3±2.9 | 95.0±4.7 | 76.3±3.4 |
| AST U/L | 41.0±4.3 | 55.3 ±4.9 | 112.7 ±4.2 | 65.3 ±3.8 | 81.0± 4.7 | 71.3 ±4.5 |
| ALP U/L | 159.7±8.6 | 172.3±5.7 | 280.0±11 | 142.3±5.8 | 194.3±8.4 | 175.7±6.3 |

Values are mean ± SEM (n=6). p≤0.05, p≤0.01, p≤0.001

## 4. DISCUSSION

Traditional medicinal plants are being investigated as a potential source of novel medications to combat the growing range of ailments and inflammatory diseases [34]. Various medicinal plants such as *Elaeis guineensis* [30] *Arnebia nobilis* [12] Ampelopsis japonica [35] and *Talinum fruticosum* [36] has been used for the assessment of wound healing and as an anti-inflammatory agents. The function of the healing process causes the restoration and replacement of damaged tissues [2] These new improvements and discoveries decreased serious complications and hospitalization regarding wound healing [37] Silver nanoparticles have acquired prominent place in disease management because of their unique properties owing to small dimension and large surface area, mechanical and thermal stability, anti-inflammatory and antimicrobial activity [38]. In the current study, the descriptions of synthesis and characterization of ABAgNPs have been published by Nazer et al. [39]. The results of this study, following the guidelines of the OECD, showed that ABaqu and ABAgNPs showed non-toxic effect on mice skin at a dosage of 1000 mg/kg body weight. Previous reported data was in agreement with recent study [40]. Hence, this study might explain the importance of investigating the safety use of ABaqu and ABAgNPs ointment on the skin. This finding demonstrates that the ABaqu and ABAgNPs ointment under study have no acute dermal toxicity. Moreover, no signs of toxicity as well as no mortality were noted during the 14 days of cage side observation. According to Hamed *et al*. [41] and Toker *et al*. [42] wound healing is a complex process that involves epithelization, inflammation, new tissue formation, and tissue remodeling. Previous literature showed that plant extracts are potential source of wound healing agents [43]. Various medicinal plants such as *Arnebia nobilis, Elaeis guineensis, Carapa guianensis* L., *Carica papaya* L. *Jasminum grandflorum* Linn, *Rubus chamaemorus, Ampelopsis radix*, and *Ampelopsis japonica* showed significant wound healing effect in treated rats [44-46, 30, 12, 35]. In current research, the wound healing effect of *A. bracteosa* was screened in Swiss albino mice and our findings agreed with the previous literature. Researchers used green synthesized silver nanoparticles to cure the wound infections and inflammation [20, 12, 47] Current results are consistent with previous investigations. Green synthesized silver nanoparticles ABAgNPs using *A. bracteosa* had significant wound healing effect due to presence of pharmacologically active phyto-constituents such as flavonol glycosides tannins, polyphenols, steroids, alkaloids, terpenoid, iridoid glycosides, reducing sugars, saponins, and flavonoids which play important role in wound healing process [48]. Younis *et al*. [49] reported showed significant antimicrobial together with wound healing abilities by biogenic AgNPs produced by Phormidium sp.

Hematological studies revealed the low level of WBC, RBC and hemoglobin in open wound (diseased) groups compared to vehicle and other treated groups, which lead to cause inflammation at the site of wound and may be occurred due to oxidative stress that increase in the production of free radicles. On the other hand, ABaqu, ABAgNPs and Nitrofuranose treated groups showed the significant increase in the hematological parameters. We can say that they involved in the reduction of free radicles due to presence of potent antioxidants (phenols and flavonoids). Some researchers have established antioxidant effects of medicinal plants which have importance to reduce the inflammation activities [50, 51, 36]. These antioxidants play an important role in wound healing. Our results are consistent with the findings of Sasidharan *et al*. [30], who reported that antioxidants of *Elaeis guineensis* has potent wound healing effect. High level of lymphocytes suppress the inflammation response during wound healing.

Histochemical analysis revealed that early regeneration of epidermis, dermis, and keratin covering in ABAgNPs and Nitrofuranose treated groups compared to vehicle group. It is confirmed that ABAgNPs had a positive effect towards epithelization, granulation tissue formation, and cell proliferation. Sasidharan *et al*. [30] also showed the early tissue regeneration in *Elaeis guineensis* treated mice. Hematoxyline and eosin staining showed the well-synthesized collagen in wound treated groups (ABaqu, ABAgNPs and Nitrofuranose), which also support the potential effect of *A. bracteosa* on fibroblast proliferation and extracellular matrix synthesis during wound healing. Alkaloids and terpenoids *A. bracteosa* may be involved in the wound healing process as reported earlier by Garg *et al*. [52]

According to John and Shobana, [36] formalin induced paw edema in mice is one of the superlative practice to display the acute anti-inflammatory effect and is a time-dependent reaction. In current study, we observed that ABaqu and ABAgNPs significantly reduced the formalin induced paw edema at 3^rd^ day of induction. This finding suggests the protective effect of green synthesized silver nanoparticles using *A. bracteosa* which has already been reported [53]. It may be due to the presence of phytochemical constituents of *Ajuga bracteosa* such as flavonoids, phenolic contents, and terpenoids which play an important role as anti-inflammatory agents. Our findings agreed with the [54-56]. Mazandarani *et al*. [57] reported that flavonoid and phenolic constituents of *N. officinale* play an important role as anti-inflammatory agent. Jang *et al*. [58] validated that AgNPs administration reduced inflammation by declining vascular endothelial growth factor, PBK, and mucous glycoprotein expression in ovalbumin-induced allergy in murine model. Wong *et al*. [59] also reported that AgNPs possessed anti-inflammatory activity in postoperative peritoneal adhesion model. Yilma *et al*. [60] showed that silver-polyvinyl pyrrolidone nanoparticles exhibited anti-inflammatory activity by reducing TNF-α.

The high level of neutrophils, lymphocytes and eosinophils were recorded in the formalin induced paw edema treated mice. Previous literature reported that leucocytes play a major role in the manifestation and development of inflammation. High level of neutrophils produced free radicals and cytokines which are also responsible for inflammation development [36]. Generation of reactive oxygen species involved in the development of inflammation [61]. Eosinophils synthesize and release lipids derivatives which involved in the inflammation development in tissues while lymphocytes cause the tissue distortion at the site of inflammation [62]. On the other hand, ABaqu and ABAgNPs treated diseased mice significantly reduced the inflammation via decreasing the population of leukocytes. We can say that *A. bracteosa* possessed strong antioxidants (phenolics and flavonoids), play critical role in the inflammation activities. The occurrence of the saponin as plant secondary metabolites are mainly responsible for anti-inflammatory property [63, 64]. Some researchers have established antioxidant effects of medicinal plants which have importance to reduce the inflammation activities [50, 51, 36] Hemoglobin and RBC play a major role in the oxygen transport. Formalin induced paw edema mice indicated the significant decrease in the RBC and Hb which leads to cause anemia. Our findings bear a resemblance to the previous study of Agnel *et al*. [62]. Similarly, the level of WBC was also increased which are responsible for the production of granulocytes stimulating factors, and involved in the development of inflammation [36]. The increased level of serum enzymes such as AST, ALT and ALP play an important role in the development of both chronic and acute inflammation. Our results are consistent with the findings of Sudharameshwari and Maheshwari [65]. The role of ABaqu and ABAgNPs against formalin induced paw edema is shown in figure 5.

**Fig. 5.**
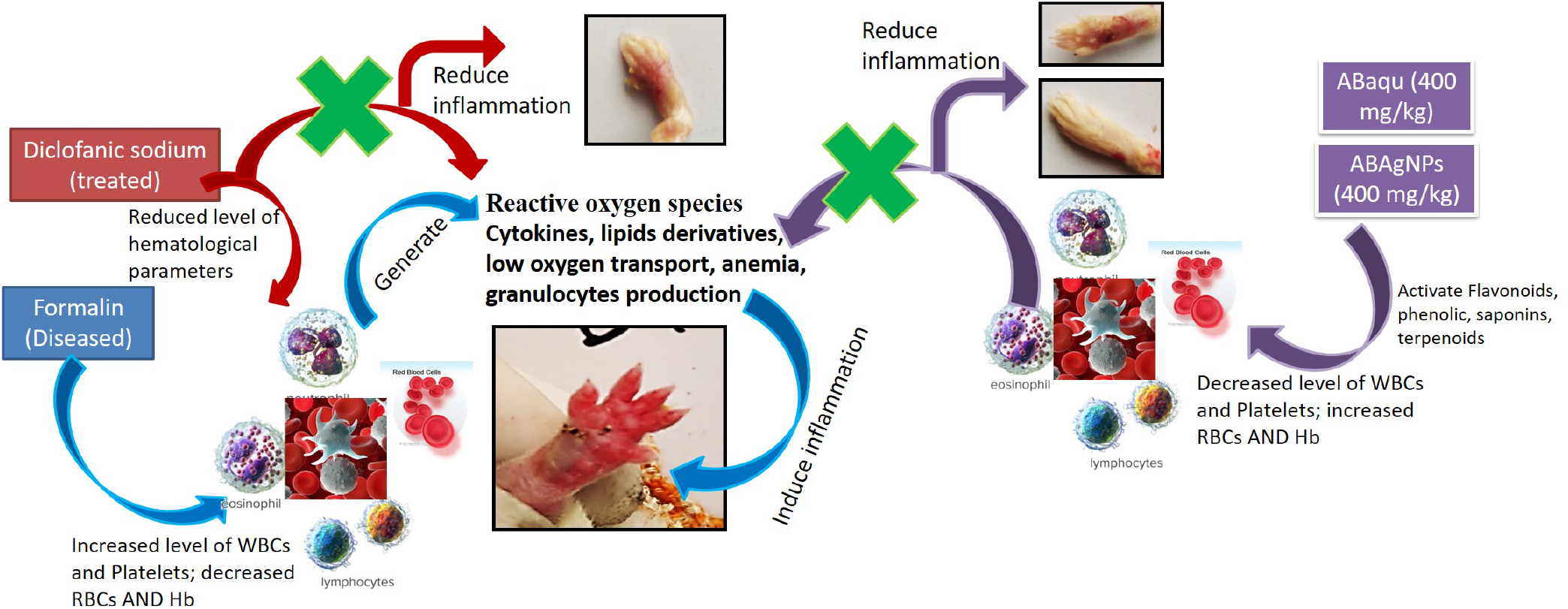
Role of *Ajuga bracteosa* conjugated silver nanoparticles and ABaqu in formalin induced paw edema Swiss albino mice.

## CONCLUSION

In conclusion, the present experimental findings of hematological, histopathological, suggests that silver nanoparticles *Ajuga bractoesa* is a promising anti-inflammatory agent in the treatment of inflammation. The phytochemical screening of the plant source contains flavonoids that confirms anti inflammatory potential of *Ajuga bractoesa*. The excision wound model showed comparable wound healing with that of standard. ABAgNPs showed faster and more effective wound healing activities than standard drug and petroleum jelly in the skin of experimentally scalded mice. Histopathological evaluation results showed better re-epithelialization, vascularization, granulation tissue formation.

## ETHICS APPROVAL AND CONSENT TO PARTICIPATE

The manuscript has been read and approved by all the authors and that the criteria for authorship have been met.

## HUMAN AND ANIMAL RIGHTS

All experiments have been designed to avoid distress, unnecessary pain and suffering to the experimental animals. All procedures were conducted in accordance with international regulations referred as Wet op de dierproeven (Article 9) of Dutch Law.

## CONSENT FOR PUBLICATION

The approval for publication of this article has been taken from all the authors.

## ACKNOWLEDGEMENTS

Authors are grateful to the Department of Physics, University of Azad Jammu and Kashmir, Muzaffarabad and National Institute for Lasers and Optronics (NILOP), Pakistan Atomic Energy Commission, Islamabad, Pakistan, for providing research facilities.

